# Evidence for selection in a prokaryote pangenome

**DOI:** 10.1101/2020.10.28.359307

**Authors:** Fiona J Whelan, Rebecca J Hall, James O McInerney

**Author notes:** **Corresponding author** Correspondence to James O. McInerney. **Contact information:** James O. McInerney, School of Life Sciences, Faculty of Medicine and Health Sciences, University of Nottingham, B41 Life Sciences Building, East Dr, Nottingham, NG7 2TQ, United Kingdom.

## Abstract

A pangenome is the complete set of genes (core and accessory) present in a phylogenetic clade. We hypothesize that a pangenome’s accessory gene content is structured and maintained by selection. To test this hypothesis, we interrogated the genomes of 40 *Pseudomonas* genomes for statistically significant coincident (i.e. co-occurring/avoiding) gene patterns. We found that 86.7% of common accessory genes are involved in ≥1 coincident relationship. Further, genes that co-occur and/or avoid each other - but are not vertically or horizontally co-inherited - are more likely to share Gene Ontology categories, are more likely to be simultaneously transcribed, and are more likely to produce interacting proteins, than would be expected by chance. These results are not due to coincident genes being adjacent to one another on the chromosome. Together, these findings suggest that the accessory genome is structured into interacting sets of genes co-selected to function together within a given strain. Given the similarity of the *Pseudomonas* pangenome with open pangenomes of other prokaryotic species, we speculate that these results are generalizable.

---

The mechanisms governing the existence of the pangenome - the totality of genes across a given set of genomes [1] - has been debated, with evidence for both neutral and selective processes [2, 3, 4]. We propose the null hypothesis that random genetic drift and gene acquisition in the absence of selection forms pangenomes. Under this hypothesis, we expect accessory gene content to have arisen as a consequence of extensive horizontal gene transfer (HGT) coupled with large effective population size, as has been argued [5]. Any observed structure in the accessory genome - including, for example, the co-occurrence of co-functional genes - would have arisen neutrally and is expected to be rare under this null model. In contrast, to observe a majority of genes overcoming the randomising effects of drift would support a rejection of the null hypothesis. Some evidence suggests that the accessory genome is under selective pressure, and that the diversity maintained is due to the selection of horizontally transferred genes which drive population differentiation and niche adaptation [2, 6]. In this case, we would expect the accessory genome to be structured into groups of genes that work well together. Similarly, we would expect genes whose interaction would be detrimental to the host to avoid being in the same genome.

To test the null hypothesis, we define gene pairs as the evolutionary unit and ask whether they are coselected across the pangenome. We focus on gene-gene association (i.e. co-occurrence) and dissociation (i.e. avoidance) patterns, collectively referred to as coincident relationships. We argue that, under the null model, we would not expect to see more coincident genes in the pangenome than would be expected by chance. In contrast, rejection of the null hypothesis would manifest as a significant proportion of the pangenome consisting of coincident gene relationships. In this case, we might further ask whether the assigned functionalities, gene expression patterns and known protein-protein interaction partners of these genes also provide evidence of co-selection. To conduct these analyses rigorously, we exclude genes that are potentially vertically or horizontally acquired together. Coincident genes that are clade-specific are likely to be coincident because they have remained within a single clade for the duration of their evolutionary history. Similarly, genes that share significant physical linkage (i.e. are co-localized on the genome) may be functionally unrelated. Removing both of these types of genes provides us with a stringent set of coincident gene pairs with which to test our hypothesis.

In this paper, we focus on the genus *Pseudomonas* as it shares properties with other well-studied open pangenomes, including persisting in a variety of niches [7] and containing comparable proportions of accessory gene content ([8, 9]; i.e. *Escherichia coli* [9, 10], *Streptococcus pneumoniae* [9, 11], *Bacillus subtilis* [9, 12]). We use coincident genes to test the null hypothesis that the microbial pangenome is maintained by drift. We identify coincident gene presence-absence patterns that deviate from random expectation, and find that 86.7% of accessory genes form ≥1 significant gene association/dissociation relationship. Co-occurring gene pairs are more likely to share functionality, be transcribed together, and to encode proteins that interact with each other more often than randomly paired accessory genes. Together, these results provide consilient lines of evidence supporting the alternative hypothesis that selection on genome content drives the evolution of the pangenome of this prokaryote.

## Results

### Species and gene distribution in the *Pseudomonas sp.* dataset

209 complete assemblies of *Pseudomonas* species were obtained from pseudomonas.com. The genomes were distributed across 40 *Pseudomonas* species, the most prevalent of which were *P. aeruginosa* (n=81), *P. putida* (n=18), *P. fluorescens* (n=15), *P. syringe* (n=13), and *P. stutzeri* (n=10) (**Supplementary Figure 1a**). 25 species were represented by a single genome within the dataset. Furthermore, a total of 22 genomes were included that do not have a species identification.

Across these 40 species, we identified a total of 96,694 orthologous gene clusters (**Supplementary Figure 1a**). Of these, only 1,365 (1.41%) were identified in ≥90% of strains (i.e. “core” genes). The mean number of genes per genome was 5,530, meaning that in a given strain, an average of 24.9% of its genes are core. PAO1 – a commonly studied *P. aeruginosa* lab strain [13] – was found to contain 5,601 genes (compared to 5,688 as annotated on Pseudomonas.com), of which 1,494 are core genes. A total of 88,792 (91.8%) genes were found in ≤15% of genomes (**Supplementary Figure 1a**). While the number of accessory genes varies across strains, the number of core genes is remarkably stable (**Supplementary Figure 1b**).

### The *Pseudomonas* pangenome contains an abundance of coincident gene relationships

Using the gene annotations provided by pseudomonas.com and gene clusters identified with Roary [14], the 96,694 orthologous gene clusters (herein referred to as gene clusters) were used to identify coincident gene relationships within the pangenome. Any gene cluster that was considered core or present in ≤5% of strains were culled from coincident analyses, leaving 13,864 gene clusters across 209 genomes for testing. From these analyses (detailed in the *Methods*), we identified a *significantly associating dataset* comprised of 293,123 cooccurring gene pairs organized into 433 connected components (**Figure 1a**). The 433 associating gene sets are well dispersed across the *Pseudomonas sp.* core gene phylogeny and none are species-specific, indicating the effect of culling lineage-dependent genes from the analysis (**Supplementary Figure 2**). Similarly, we determined the *significantly dissociative dataset* which contains 421,080 dissociative gene pairs organized into 13 connected components (**Figure 1b**).

**Figure 1:**
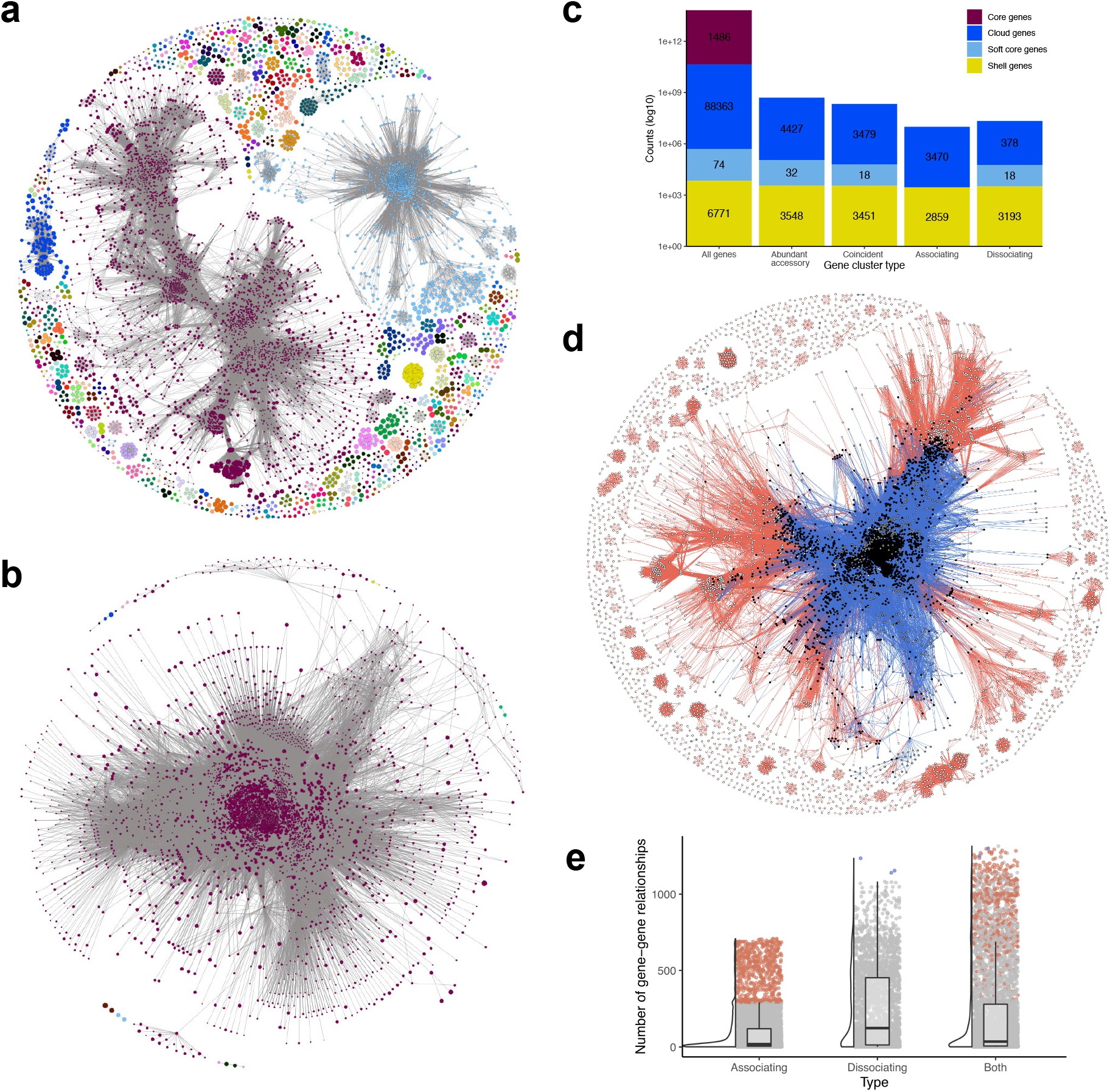
Network of coincident relationships in the *Pseudomonas sp.* accessory pangenome. Relationships between significantly associating (**a**) and dissociating (**b**) gene pairs are shown as gene-gene networks. Only nodes with a D*≥*-0.4 (i.e. sufficiently lineage-independent) are displayed. Nodes (i.e. gene clusters) are connected to other nodes if-and-only-if there is a significant coincident relationship between them. Nodes are coloured by the connected component which they below to; in other words, nodes are coloured by significantly coincident gene sets. The size of the node is proportional to the D-value of the gene cluster (the larger the node, the more lineage-independent the gene is); the thickness of the edge is reversely proportional to the p-value associated with the coincident relationship. **c.** Of the abundant accessory subset of all lineage-independent genes within the pangenome, 86.7% are involved in coincident relationships. **d.** A gene-gene network of all lineage-independent coincident gene relationships. Edges are coloured by association (**red**) and dissociation (**blue**) relationships. Genes which form both association and dissociation relationships are represented by **black** nodes, genes which only associate by **white**, and genes which only dissociate by **gray**. **e.** The distribution of gene-gene relationships across genes. Boxplots display the first and third quartiles, with a horizontal line to indicate the median, and whiskers extend to 1.5 times the interquartile range. Associating and dissociating “hub” genes are coloured.

**Figure 2:**
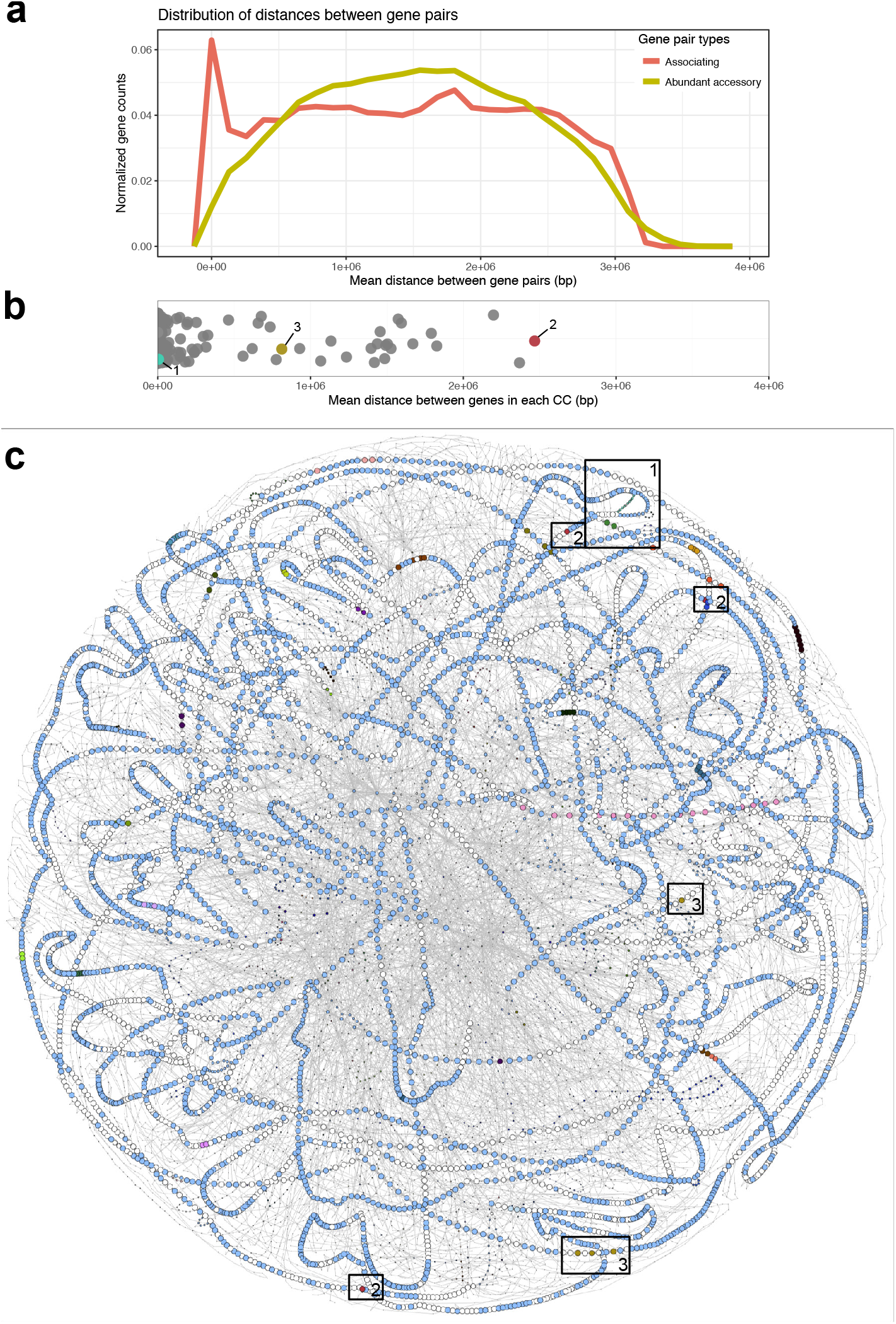
Co-localization amongst associating gene pairs. **a.** Associating genes are more likely to be colocalized than are randomly assigned abundant accessory gene pairs on *Pseudomonas sp.* chromosomes. **b.** 26% of all sets of associating genes (i.e. connected components of genes which share co-occurrence patterns) do not share significant physical linkage as defined by the mean distance between all genes within a gene set. Coloured gene sets correspond to labelled boxes in part C. **c.** The pangenome graph of the *P. aeruginosa* subset of the *Pseudomonas* dataset. The pangenome graph of the full dataset is available in **Supplementary Figure 5**. Labelled boxes show examples of gene association clusters that are co-localized (box 1, turquoise genes), are not co-localized (boxes 2, red genes), and have variable levels of genetic distance (boxes 3, green genes). For visibility, cloud genes are not shown.

Of the 13,864 accessory gene clusters identified in ≥5% of *Pseudomonas* strains (i.e. the abundant accessory genes tested by Coinfinder [21]), 8,007 (57.7%) were lineage-independent (see *Methods*, **Supplementary Figure 3**). Of these 8,007 clusters, 6,329 and 3,589 formed associating and dissociating relationships, respectively (**Figure 1c**). Accounting for the genes involved in both types of relationships, a surprising 6,948 (86.7%) of abundant lineage-independent accessory genes were involved in ≥1 coincident relationship. While gene dissociations were identified across all three non-core gene categories, gene associations were only identified in the two more rare gene categories (Cloud and Shell genes; **Figure 1c**). Similar results were found when both lineage-independent and -dependent genes were considered (**Supplementary Figure 4a**).

Of the 6,329 genes forming coincident relationships identified, 2,970 (46.9%) are involved in both association and dissociation relationships, meaning that they both co-occur with, and avoid other genes in the pangenome (**Figure 1d; black nodes**). These 2,970 dual-relationship genes account for 268,647 (91.6%) of all gene-gene associations and 418,698 (99.4%) of all gene-gene dissociations (**Figure 1d**). That is to say that almost half of the coincident genes account for the majority of coincident gene relationships. On average, associating genes form relationships with 94 other genes (**Figure 1e**). However, the distribution is uneven, with 24.3% of genes forming fewer than five connections to other genes (1,542 genes < the 25th percentile; **Figure 1e**). The 624 association hubs (i.e. genes with *>*1.5x the upper interquartile range) each have ≥290 gene associations and account for 50.8% number of dissociation hub genes (n=3) that each form ≥1,110 gene dissociation relationships. Among the associating and dissociating hub genes are a diversity of functions including transcriptional regulators, transporter subunits, metabolic enzymes, and an abundance of hypothetical proteins. Interestingly, for those genes that were found to have both types of coincident relationships, no gene acts as both an associating and dissociating hub (**Figure 1e**). The number of hub genes increase when lineage-dependent genes are included in these analyses (**Supplementary Figure 4b**).

### Co-localization of coincident genes

HGT and differential gene loss are the main contributing factors to pangenome formation [15]. If functionally related gene pairs are found in close proximity on a genome, then they may have been acquired in a single HGT event, and their co-occurrence pattern might be a consequence of the HGT process, and not a consequence of natural selection. However, many known protein interactions occur between genes that are dispersed across the genome (for e.g. proteins produced by genes *crr* and *ptsG* form the the EII complex in enteric bacteria and are not in close proximity on the genome [16]). To explore whether co-localization and the simultaneous transfer of genes is responsible for gene association relationships in the pseudomonads, we compared the mean syntenic distance of associating genes, versus the mean syntenic distance of abundant accessory gene pairs chosen at random. The average chromosome length across the dataset is 6.2 Mbps; which, in addition to the chromosome being circular, means that the furthest away two genes could be from each other is ~3.1 Mbps. The mean distance between randomly paired abundant accessory genes is bell-shaped which fits our expectation of randomly dispersed genes. In contrast, associating gene pairs more often share significant localization (**Figure 2a**); however, only 8.6% of all co-occurring gene pairs have a mean distance of <150kbp. This suggests that a proportion of, but not all, gene-gene co-occurrence is due to co-localized genes.

In order to ask whether the co-localization patterns of gene pairs generalize to that of gene sets, we next considered gene associations in terms of their connected component (i.e. associating gene set; **Figure 1a**). We observe 41 gene sets (26%) that are composed of pairs of genes with a mean pairwise distance of ≤150 kbp (**Figure 2b**). We used PPanGGOLiN [17] to generate pangenome graphs of *Pseudomonas sp.* (**Supplementary Figure 5**) and the *P. aeruginosa* subset (**Figure 2c**) to visualize the genomic context of co-localized gene sets. For example, the *P. aeruginosa* pangenome graph includes a set of neighbouring co-occurring genes associated with flagellar assembly (**Figure 2c, box 1**). Interestingly, this path in the pangenome graph bypasses a set of 16 genes which also show homology to flagellar assembly genes (**Supplementary Table 1**). A given genome may contain one but not both of these sets of genes, indicating possible redundancy of this function within the pangenome. We also observe gene sets that share very little physical linkage, such as a set of three unnamed genes involved in outer membrane permeability (**Figure 2c, box 2; Supplementary Table 1**). Still, other gene sets have mixed levels of co-localization amongst their membership. For example, a subset of *P. aeruginosa* strains contain three neighbouring genes that co-occur with a fourth gene sharing no physical linkage with the other three (**Figure 2c, box 3**); these four genes likely co-occur because they all function as components of the methionine salvage pathway (**Supplementary Figure 6**, **Supplementary Table 1**).

### Coincident genes share functionality

The association (or dissociation) of genes alone does not infer a biological interaction between them (i.e. correlation does not infer causation; [18]). In order to reject the null hypothesis that the accessory genome is governed by random genetic drift, we would expect that coincident genes would be more likely to act together - for example, towards a shared functional goal - for the benefit of the host. Using Gene ontology (GO) annotations as a proxy for gene functionality, we calculated the functional overlap of each coincident gene pair in comparison to randomly paired abundant accessory genes (**Figure 3a**). We identified a greater overlap in GO annotations between coincident gene pairs then randomly paired accessory genes. Specifically, 71.1% of associating and 69.4% of dissociating gene pairs shared GO annotations when compared to only 50.6 (±0.1)% of randomly paired accessory genes (**Figure 3a**). This indicates that coincident genes share function with each other more often than would be expected by chance. The percentage of shared GO annotations amongst associating genes increased to 74% when only non-syntenic genes were considered (**Supplementary Figure 7**). Given these results, we calculated whether particular GO terms were more likely to share annotation in a coincident gene pair compared to the expected term-sharing frequency (**Figure 3b**). 150 GO terms were found to be overrepresented in gene-gene associations, including pilus assembly (GO:0009297; p=1.41e-05), type II protein secretion system complex (GO:0015627; p=1.35e-08), and antibiotic biosynthetic process (GO:0017000; p=4.84e-10) (**Figure 3b red points, Supplementary Table 2**). In contrast, 60 GO terms were overrepresented in dissociation relationships, including ATP-binding cassette (ABC) transporter complex (GO:0043190; p=4.96e-52), and drug transmembrane transport (GO:0006855; p=2.16e-07) (**Figure 3b blue points, Supplementary Table 2**).

**Figure 3:**
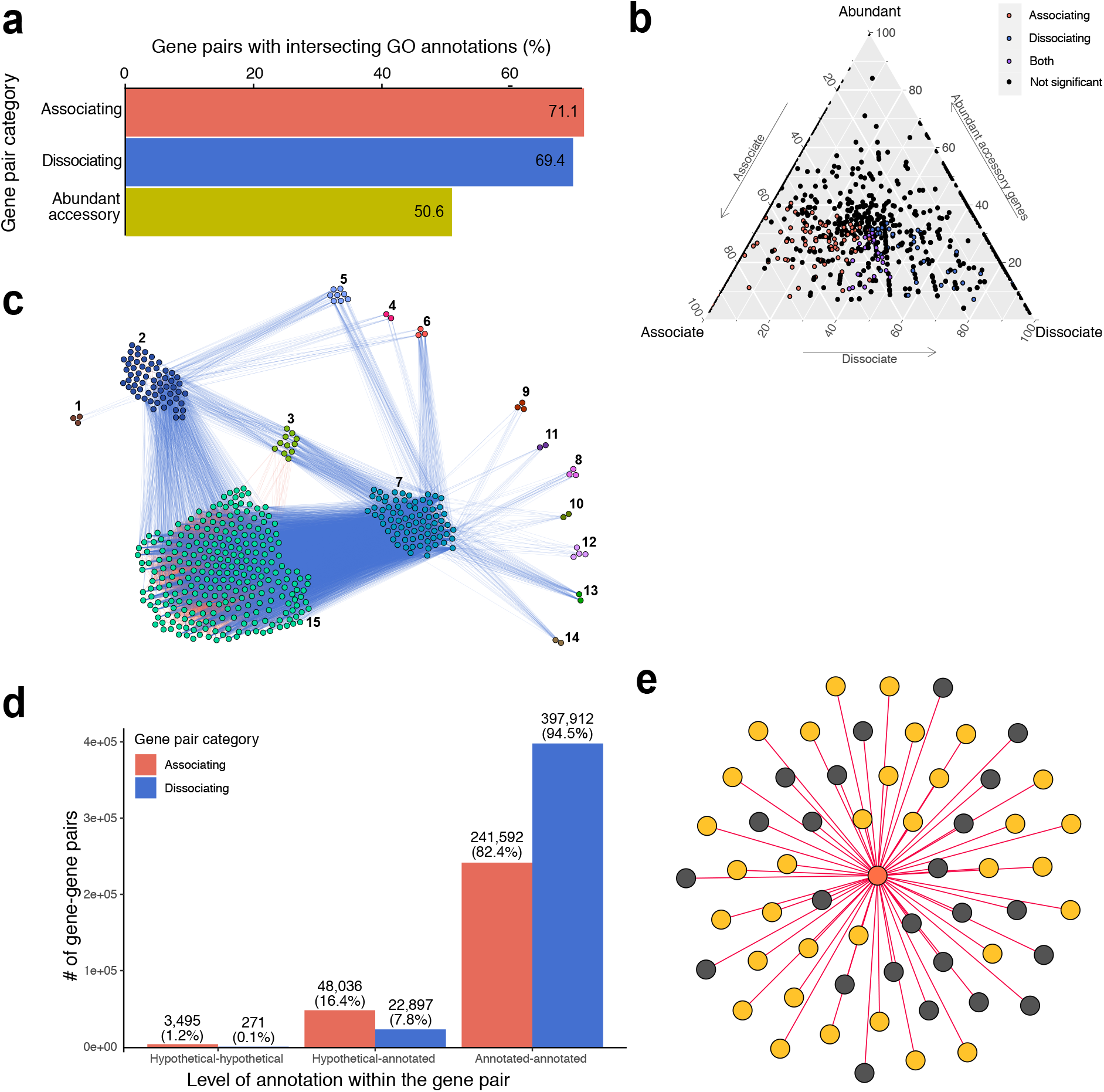
Coincident (associating and dissociating) gene pairs have more overlapping GO term annotations when compared with random gene pairs. **a.** 71.1% of associating gene pairs share the same GO annotations compared with 50.6 (*±* 0.1)% of randomly paired genes. **b.** Triangular plots of GO term annotation within coincident gene space. Each GO term is represented by a point whose location is determined by how frequently genes with that term are found in the associating, dissociating, and random gene pair categories. GO terms which are significantly overrepresented in any category are coloured **c.** Coincident gene relationships for genes annotated with transmembrane transporter activity (GO:0022857). Edges are coloured by the type of interaction (associating, **red**; dissociating, **blue**). A Figure showing only the associating edges is provided in **Supplementary Figure 8a**. **d.** The proportion of coincident gene pair relationships which exist between annotated and hypothetical genes. **e.** A network of gene (node) association relationships (edges) depicting the associations of *ampC* (**orange**) with hypothetical (**gray**) and annotated (**yellow**) genes.

A subset of GO annotations was enriched in both associating and dissociating gene pairs (**Figure 3b purple points; Supplementary Table 2**). This appears counterintuitive, but may correspond to, for example, two multi-gene functional units that dissociate from one another but whose genes within the unit strongly associate with each other. For example, gene pairs annotated with transmembrane transporter activity (GO:0022857) were enriched in association (p=8.39e-06) and dissociation gene relationships (p=3.01e-28; **Figure 3c**). While some genes formed independent co-occurring cliques or solitary dissociation patterns (not shown), the majority of genes clustered into groups of associating genes (**Supplementary Figure 8a**) that dissociated from each other (**Figure 3c**). Some of these cluster avoidance patterns appear to be largely due to species boundaries (e.g. clusters 7 and 15; **Supplementary Figure 8b**) but most are independent of phylogeny and syntenic relationships (**Supplementary Figure 8bc**). Although many of these genes are hypothetical or only loosely annotated, there are, for example, genes for an efflux pump (Resistancenodulation-division (RND) family transporters) in cluster 2 that dissociate from genes for a different efflux pump (glutathione-regulated potassium-efflux system protein, KefB) in cluster 3 (**Supplementary Table 3**), indicating a possible example of functional redundancy or niche partitioning within this system.

The above calculations of intersecting GO annotations rely on known gene information. While *Pseudomonas sp.* is a well-studied genus with well-annotated genomes, many of the identified coincident gene pairs involve interactions between hypothetical proteins or genes without a known GO association. 51,531 (17.6%) and 23,168 (7.9%) of the associating and dissociating gene pairs, respectively, involve at least one hypothetical gene (**Figure 3d**). Specifically, 95% of coincident gene pairs involving hypothetical genes are between hypothetical and annotated genes. Given our finding that many annotated coincident gene pairs share function, coincident relationships between hypothetical and annotated genes can help us generate hypotheses concerning the role these hypothetical proteins play in the *Pseudomonas sp.* pangenome. A subset of GO terms was found to be statistically more likely to be coincident with hypothetical genes when compared to the annotated coincident gene pairs (**Supplementary Table 4**). For example, the “beta-lactamase activity” (p=1.86e-06; GO:0008800) GO annotation was assigned to two genes that collectively associated with 120 annotated and 33 hypothetical genes. In particular, 42% of the genes that associate with an *ampC* homolog (most closely related to PDC-8 [19]) were annotated as hypothetical proteins, and only seven had a gene name annotation in ≥1 genome (**Figure 3e, Supplementary Table 5**). This gene association cluster (including *ampC*) is present in ≥4 *Pseudomonas* species (4 named, 6 unnamed strains), and does not share considerable co-localization across the pangenome (**Supplementary Figure 9**). Similar investigations of the remaining hypothetical-annotated gene pairs may yield further hypotheses concerning the role of hypothetical proteins in this pangenome.

### Gene co-occurrence is associated with co-transcription and protein-protein interactions

Using publicly available RNA-Seq transcription data, we examined how often associating gene pairs were transcribed together compared to randomly paired accessory genes. Due to limitations on the availability of good quality publicly available gene transcription data, we restricted our analysis to *P. aeruginosa* (81 of 209 genomes). Across the *P. aeruginosa* pangenome, we calculated the frequencies with which a given gene pair was transcribed together compared to that of only one of the two genes in a pair. We report this ratio of gene expression, such that a ratio of 1.0 indicates that - across the *P. aeruginosa* pangenome - it is as likely to see both genes transcribed together as it is for only one of the pair to be transcribed (**Figure 4a**). Across samples and experiments, associating gene pairs were more often co-transcribed than were randomly paired abundant accessory genes (**Figure 4a**), indicating a possible shared function or interaction between these genes. This result holds when only non-syntenic gene associations are considered (**Supplementary Figure 10**). Similar analyses of co-transcription could not be performed on the dissociating gene pairs as these pairs are not present within the same genomes.

**Figure 4:**
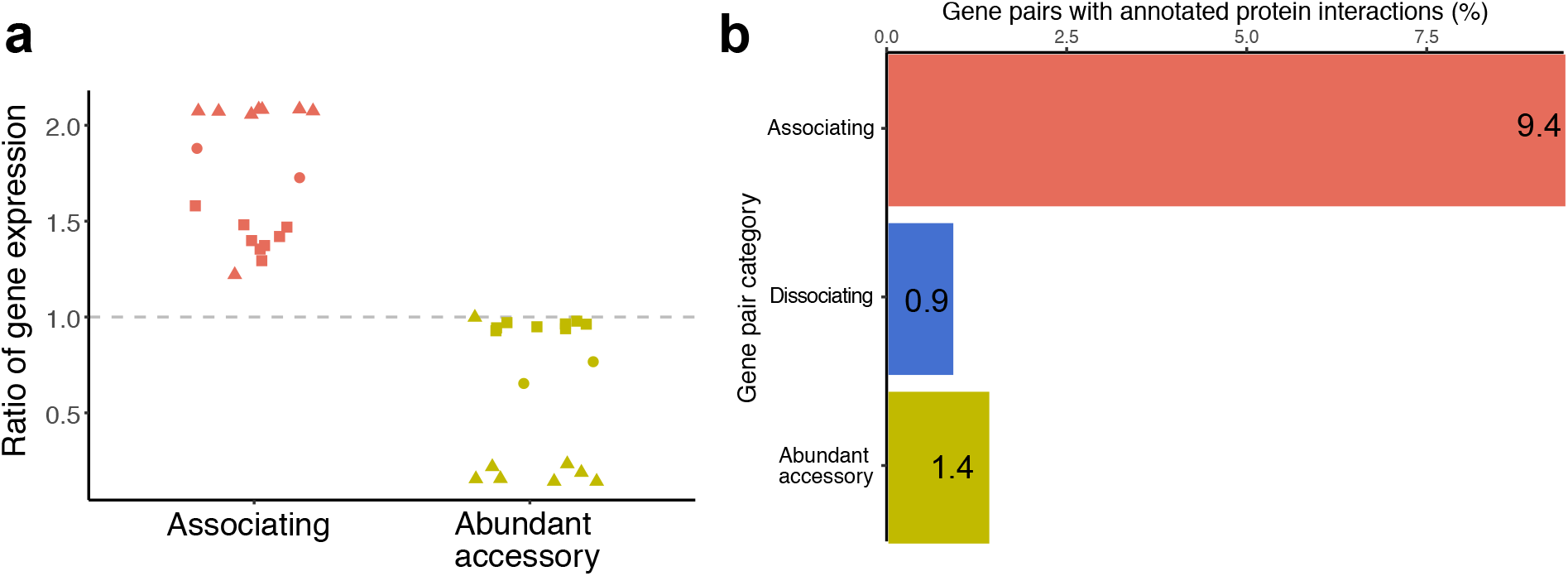
Associating genes are more likely to be co-transcribed. **a.** The ratio of gene expression between associating gene pairs and random abundant accessory gene pairs. The ratio is calculated as the proportion of times that both genes in a gene pair are consistently co-transcribed across *P. aeruginosa* genomes versus the proportion of times that only one of the two genes is transcribed. Symbols represent different publicly-available RNA-Seq experimental projects. **b.** Protein-protein interaction pairs as compared to the STRING database indicate more interactions in the associating gene pairs compared to the dissociating and random gene-gene data. 100 replicates of randomly paired genes were used to obtain a mean of 1.4 (*±*0.03)%.

Given the rate of co-transcription of associating genes, we asked how often coincident genes are involved in known protein-protein interactions. Using the STRING database [20], we first identified the number of protein-protein interactions between randomly paired accessory genes as 1.4 (±0.03)%. This percentage may seem low; however, we expect that documented protein-protein interactions are more likely to involve well-studied, abundant (likely core), house-keeping proteins, or those which share evolutionary histories with each other, which are precisely the genes which are excluded in our analyses of linkage-independent accessory genes. However, we identified protein-protein interactions between 9.4% of associating gene pairs (11.4% of all annotated associating pairs; **Figure 4b**). These data represent 2.5% of all known protein-protein interactions within *P. aeruginosa*; that is to say that - even when excluding core or vertically inherited genes - associating gene relationships recapitulates a percentage of all known protein interactions in this species. As expected, evidence of interactions between dissociating genes were identified at a rate less than randomly paired genes (**Figure 4b**).

## Discussion

We recently developed a novel method for the identification of coincident gene presence-absence patterns within pangenomes [21]. Here, we applied this approach to 209 publicly available *Pseudomonas sp.* genomes to test the null hypothesis that pangenome gene content is determined by random genetic drift. Across the dataset, 86.7% of lineage-independent, abundant accessory genes consistently associated with, or dissociated from, at least one other gene in the pangenome. This represents a lot more genetic structure within the accessory genome than we would expect if neutral processes were driving pangenome formation. We found that these gene pairs share functional annotations, are co-transcribed, and produce proteins that interact with each other more often than expected when compared to randomly paired abundant accessory genes. These findings were independent of genes which have a significant phylogenetic signal (i.e. are lineage-dependent or are predominantly vertically transmitted) and was also the case when co-localized genes were excluded. The fact that we found statistically significant associations between non-syntenic genes is strong evidence allowing us to reject the null hypothesis because it identifies genes that share functionality despite being dispersed in the genome. Together, these data suggest that the assemblage of accessory genes in this pangenome does not conform to the expectation that random genetic drift has dominated its evolutionary history. Instead, we propose the alternative hypothesis that the accessory pangenome is governed by selection. This work has implications for our understanding of prokaryote pangenomes as a whole.

We were very careful in our interpretation of these results to refer to gene-gene co-occurrences as “associations” and not “interactions”. Although such a high-throughput examination of gene-gene co-occurrence relationships in pangenomes may be rare [22, 23, 24], there is a century of literature on species-species cooccurrence patterns [18, 25, 26, 27, 28]. In this research, it has been explicitly shown that in at least some cases, species-species co-occurrence does not necessarily imply species-species ecological interactions. In their recent Perspectives article, Blanchet *et al.* present seven arguments for why ecological interaction should not be assumed from co-occurrence data [18]. Although some of these arguments are species-specific, many apply to gene-gene data as well. For example, the authors argue that in some cases, species occurrences depend on the environment, and what appears as a species-species co-occurrence pattern may actually be an indirect interaction of both species with their environment [18]; similarly, *geneA* and *geneB* may co-occur due to a preference for a shared abiotic factor - environment, nutrient, metabolite etc. - instead of indicating a direct gene-to-gene interaction. We suggest that further *in vitro* investigations of gene pairs could help clarify these levels of interactions. Further, the methodology used here - the identification of coincident gene relationships based on statistically similar or dissimilar gene presence/absence patterns - will not identify all associations in the pangenome. For example, asymmetrical dependencies will have been missed; in the case where *geneA* relies on *geneB* for an interaction but not vice versa, we would expect to see *geneA* present only in the presence of *geneB*, but that *geneB* could be present without *geneA* in a given genome. So called “event horizon genes” or those genes whose presence “leads the way” for the introduction of many other genes [29], will also not be identified by use of the Coinfinder software. Because these gene-gene patterns are hard to distinguish from random presence/absence patterns, their influence on the structure of the pangenome will be harder to determine.

With this caveat in mind, we sought to provide evidence for the possibility that a sizable subset of the gene-gene associations within the *Pseudomonas* pangenome may be due to direct interactions. The fact that many associating gene pairs tend to neighbour each other indicates this potential. Neighbouring genes often assemble into sets of co-transcribed genes which either physically interact to form protein complexes (e.g. *manXYZ* [30]) or act as part of a shared metabolic pathway (e.g. the *lac* operon [31]). However, many coincident genes which were not co-localized had overlapping functionality. These genes could still directly interact, although could also indicate a response to a shared abiotic factor (for e.g. genes present in response to a particular environmental niche). On the other hand, genes with shared functionalities which actively avoid each other would seem to suggest a more directed type of interaction. Either way, evidence for interactions at the protein level clearly indicate direct gene-gene interactions in the accessory pangenome.

One of the inspirations for this work was the recent suggestion that one way of better elucidating whether the pangenome is evolving neutrally or adaptively was to focus on the gene as the evolutionary unit [3]. Examining gene-gene relationships, as we have done here, is not the only gene-focused approach to understanding the evolutionary pressures present on prokaryote pangenomes. For example, analyses could be conducted to determine whether accessory genes are under selective pressures. Further, gene knockout and long-term evolutionary experiments could be combined to determine the effect of individual genes on the pangenome. We propose these results concerning gene-gene coincident relationships as one line of evidence for testing hypotheses of selective pressures on the accessory genome. We encourage further work in these areas to be contributed to this debate.

We focused our analysis on *Pseudomonas sp.* due to its diverse, well-studied pangenome [8, 32, 33, 34, 35], well-annotated genomes [36], and generalizability to other prokaryotic open pangenomes in terms of core-to-accessory gene ratios, and multiple environmental niches. Our results suggest genetic structure within this pangenome, and we hope that additional research, using different methodologies and pangenomes, will help further these findings.

## Methods

### Sequence acquisition & pangenome analysis

Genome annotations were retrieved from pseudomonas.com in GFF3 format [36] on 1 March 2019 and include 209 complete genome assemblies. Despite the availability of thousands of draft genomes, we restricted our study to completely assembled and curated strains, due to recent work suggesting that the quality of genome assembly can greatly impact predicted pangenome quality and size [37]. Genes were clustered into gene families using Roary 3.12.0 [14] with a 70% BLASTP percentage identity cutoff. Definitions of core (90%≤x≤100%), soft core (89%≤x<90%), shell (15%≤x<89%), and cloud (*x*<15%) genes are as in Roary. All core genes (present in ≥90% of *Pseudomonas* genomes) were individually aligned using MAFFT v7.310 [38], the alignments concatenated, and curated using Gblocks ([39]; parameters as in [40], specifically allow gap positions = half, minimum length of block = 2). A core gene phylogeny was constructed from this curated and concatenated alignment using IQ-TREE [41] using the GTR+I+G substitution model (as justified in [42]). A total of 19 genome annotations contained plasmids which were not considered in these analyses.

### Evaluation of gene coincident relationships

Coincident relationships between gene pairs were determined using Coinfinder [21]. Briefly, for each pair of genes in the input accessory genome, Coinfinder examines their presence/absence patterns to determine if they represent a coincident relationship (i.e. if they co-occur or avoid each other across the pangenome more often than expected by chance). Statistically significant coincident gene pairs were determined by Coinfinder *via* a Bonferroni-corrected binomial exact test statistic, and the lineage dependence of each gene was calculated using a previously established phylogenetic measure of binary traits (D; [43]). Coinfinder was run with upper- and lower-filtering gene abundance thresholds of 90% and 5%, respectfully. A threshold of D≥−0.4 was used based on the frequency of genes and their distribution across species in the core gene phylogeny (**Supplementary Figure 3**). The resulting associating and dissociating networks were visualized using Gephi [44]. Hub genes were defined as those with more gene-gene relationships than 1.5 times the upper interquartile range.

In order to determine whether coincident gene pairs were more likely to share functional annotations, gene expression patterns, or protein-protein interactions (see below), we compared these results against the null model by generating random abundant accessory gene pairs. To do so, accessory genes that were included in the Coinfinder analysis (i.e. were between 5-90% abundance with D≥−0.4) were paired at random to match the mean number of associating/dissociating gene pairs (n=357,102) in 100 replicates (herein referred to as random abundant accessory gene pairs). This was accomplished by creating a list of all possible paired combinations of abundant accessory gene pairs and creating n=100 random permutations of the list to a length of 357,102. The specific use of these random abundant accessory gene pairs is outlined in the following Methods sections.

### Gene co-localization and pangenome structure analysis

The physical linkage between genes in a gene pair was determined both for associating, and for random abundant accessory gene pairs. For a given gene pair, the physical distance between *geneA* and *geneB* was calculated for each genome for which both *geneA* and *geneB* reside. (For this reason, distance information could not be calculated for dissociating gene pairs.) From these *geneA-geneB* distances for each genome, a mean distance was computed and plotted. In analyses of non-syntenic genes, only those gene pairs separated by a mean distance of ≥150 kbp were considered.

A pangenome graph was created with PPanGGOLiN [17]. In order to maintain consistency with the gene cluster information used throughout this study, PPanGGOLiN was provided with the gene clusters as determined by Roary. A Python script was used to redefine nodes in the pangenome graph to remain consistent with the definitions of core, soft core, shell, and cloud that are used by Roary. The nodes of the resulting graph were recoloured to represent the associating gene sets as determined by Coinfinder. The network was visualized in Gephi [44]. KEGG was used to investigate metabolic pathways [45].

### Functional annotations of coincident genes

Gene ontology (GO) term annotations for each of the 209 genomes were collected from pseudomonas.com on 22 March 2019. A minimum of one matching GO term annotation was necessary to consider a gene pair as having overlapping function. Overlapping annotations were determined by examining only those gene pairs for which both genes had a GO term annotation. After removing gene pairs for which GO term annotations were missing for one or both genes, a total of 246,637 (84.1%) associating, and 379,439 (90.11%) dissociating gene pairs remained. These were compared to 100 replicates of randomly paired abundant accessory genes as described above. Bonferroni-corrected binomial tests (computed in R [46]) were used to determine which GO terms were under- or over-represented in the coincident gene pairs when compared to the random abundant accessory gene pairs.

Separately, GO terms which were significantly associated with genes of hypothetical function was determined. Genes were defined as hypothetical if every instance of the gene across all genomes in which it was found were annotated as “hypothetical protein”. Bonferroni-corrected binomial tests were used to determine GO terms over-represented in gene pairs involving an annotated and hypothetical gene. Sub-networks of specific gene-gene interaction pairs were displayed using Gephi [44].

### Gene expression analysis

Short read archive (SRA) transcript data from the following *P. aeruginosa* RNA-Seq experiments (paired-end reads with a range of 4,450,537 - 41,817,822 reads per sample) were used to test co-transcription levels of gene-gene pairs: SRP163899 (n=2 samples), SRP215630 (n=9), and SRP191772 (n=8; [47]). The reads from each RNA-Seq sample were mapped using Bowtie2 [48] to the gene content of the *P. aeruginosa* genomes in the dataset (n=81). In a given genome, a gene was considered transcribed if ≥85% of the gene’s length was covered by ≥2 reads. Across the dataset, a gene cluster was considered transcribed if it was transcribed in ≥75% of the genomes in which it was present. The ratio of gene expression is the ratio of gene cluster pairs which are co-transcribed versus those in which only one of the two genes were transcribed. Therefore, a ratio of 1.0 would mean that, across all *P. aeruginosa* genomes, paired genes are just as likely to be co-transcribed as for exclusively one of the two genes to be transcribed; a ratio of 2.0 would mean that paired genes are twice as likely to be transcribed together across the pangenome.

### Protein interaction analysis

The STRING database [20] was used to identify whether the protein products of associating, dissociating, and random abundant accessory gene pairs interact with each other. The protein network data and associated FASTA sequences for *P. aeruginosa* were obtained from https://string-db.org. The FASTA sequences for the proteins in this network were assembled into a BLAST database to map homologous gene clusters to the IDs in the STRING protein network, with the criteria of ≥85% coverage and ≥90% sequence identity. Calculations of the coincident gene pairs were compared to 100 replicates of randomly paired abundant accessory gene pairs as described above.

## Data Availability

All raw data, including genome and gene identifiers, used in this work is available as a SQL Schema from github.com/fwhelan/pseudomonas-manuscript including maps between genomes, genes, gene clusters, and GO term annotations. An R markdown file, pseudomonas manscript.Rmd, available at github.com/ fwhelan/pseudomonas-manuscript details how each Figure was generated from the available raw data.

## Code Availability

The set of Python scripts and SQL queries used to generate data matrices, and an R Markdown file of the R code used to generate all Figures are available from github.com/fwhelan/pseudomonas-manuscript.

## Acknowledgements

This research was funded by a Marie Skłodowska-Curie Individual Fellowship (GA no. 793818) awarded to FJW and BBSRC Responsive Mode Grant BB/N018044/1 awarded to JMcI. We would like to thank the tireless efforts of The *Pseudomonas* Genome Database and their funders. We acknowledge critical intellectual conversations with P. Mulhair and M.R. Domingo-Sananes.

## Author Contributions

FJW is the primary author of this prepared manuscript. FJW collected, processed, and analysed all data. RJH provided key intellectual insights and Figures for all metabolic pathways considered within. FJW, and JOM conceptualized the experimental outline. FJW conducted all data analyses and wrote this manuscript. All authors edited and approved the manuscript.

