## Supplemental Information for "Evidence for selection in a prokaryote pangenome"

### Supplementary Information

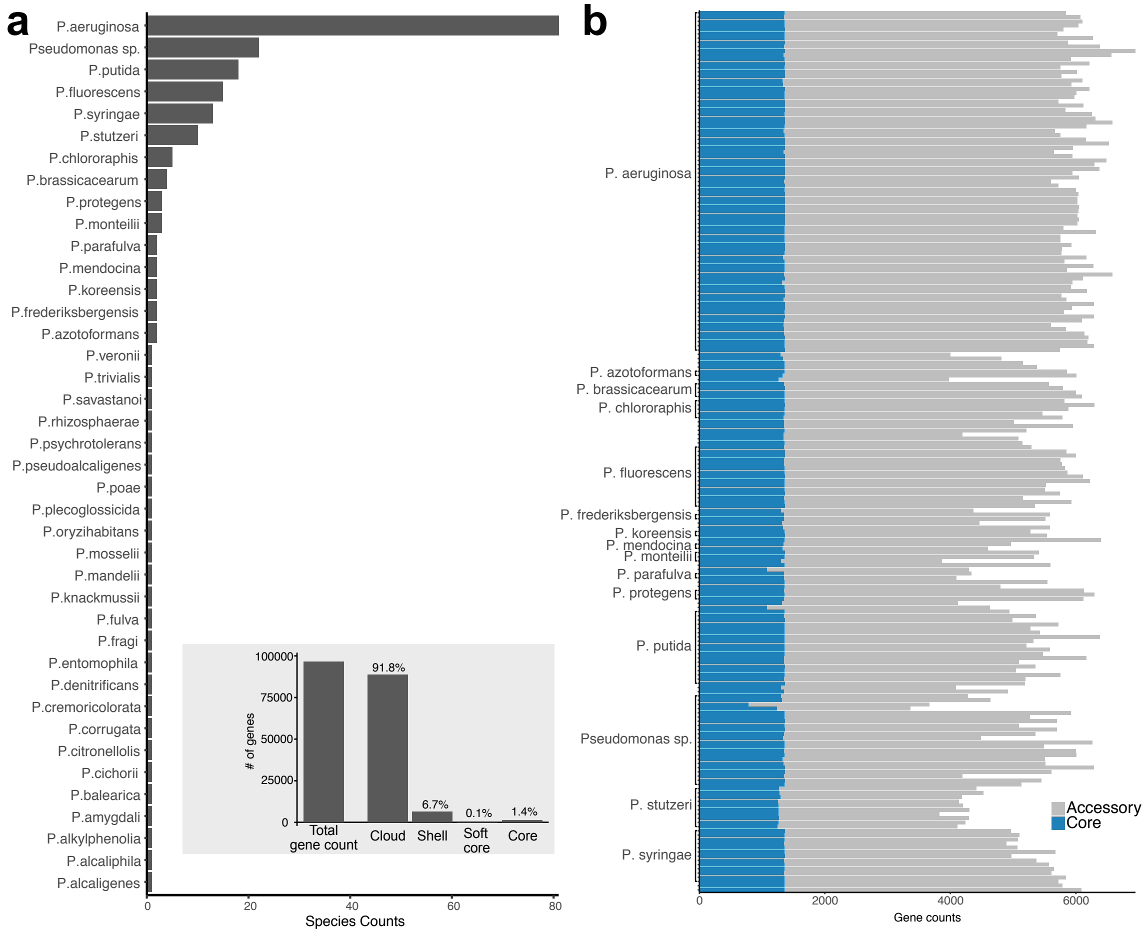


Supplementary Figure 1: **Distribution of *Pseudomonas* species and genes in the dataset. a.** The number of genomes per species present in the dataset (n=209). **a, inset.** The total gene cluster counts (n=96,694) across all strains broken down into the following categories: Cloud genes (present in 0% ≤ strains < 15%), Shell genes (15% ≤ strains < 89%), Soft core genes (89% ≤ strains < 90%), and Core genes (90% ≤ strains ≤ 100%). Collectively, cloud, shell, and soft core genes are referred to as accessory genes. **b.** The distribution of core and accessory genes across all strains.

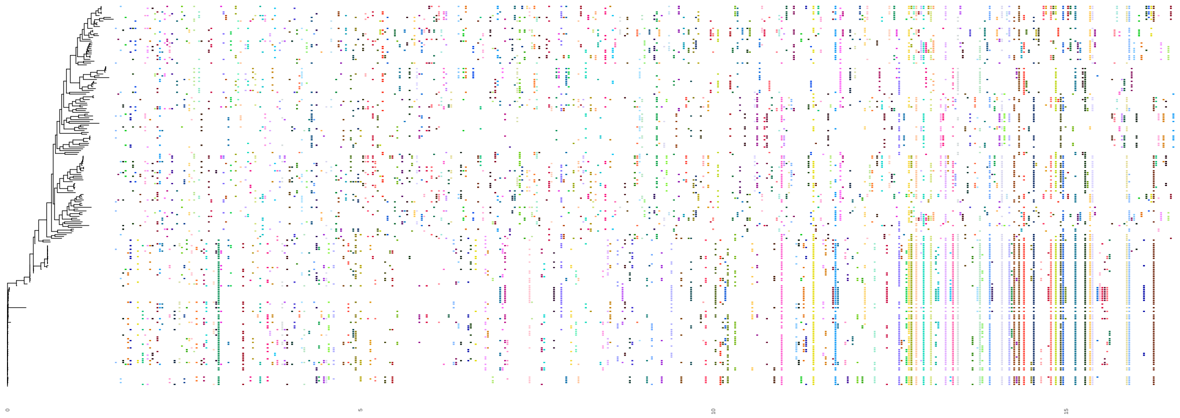

Supplementary Figure 2: **Mean presence of the genes comprising each connected component in the association dataset.** Each connected component (i.e. set of genes which associate with each other) is well dispersed across the core gene phylogeny indicating that vertically inherited, or lineage-dependent genes, have been successfully culled from the dataset. Each connected component is indicated by a single, randomly coloured column in the heatmap.

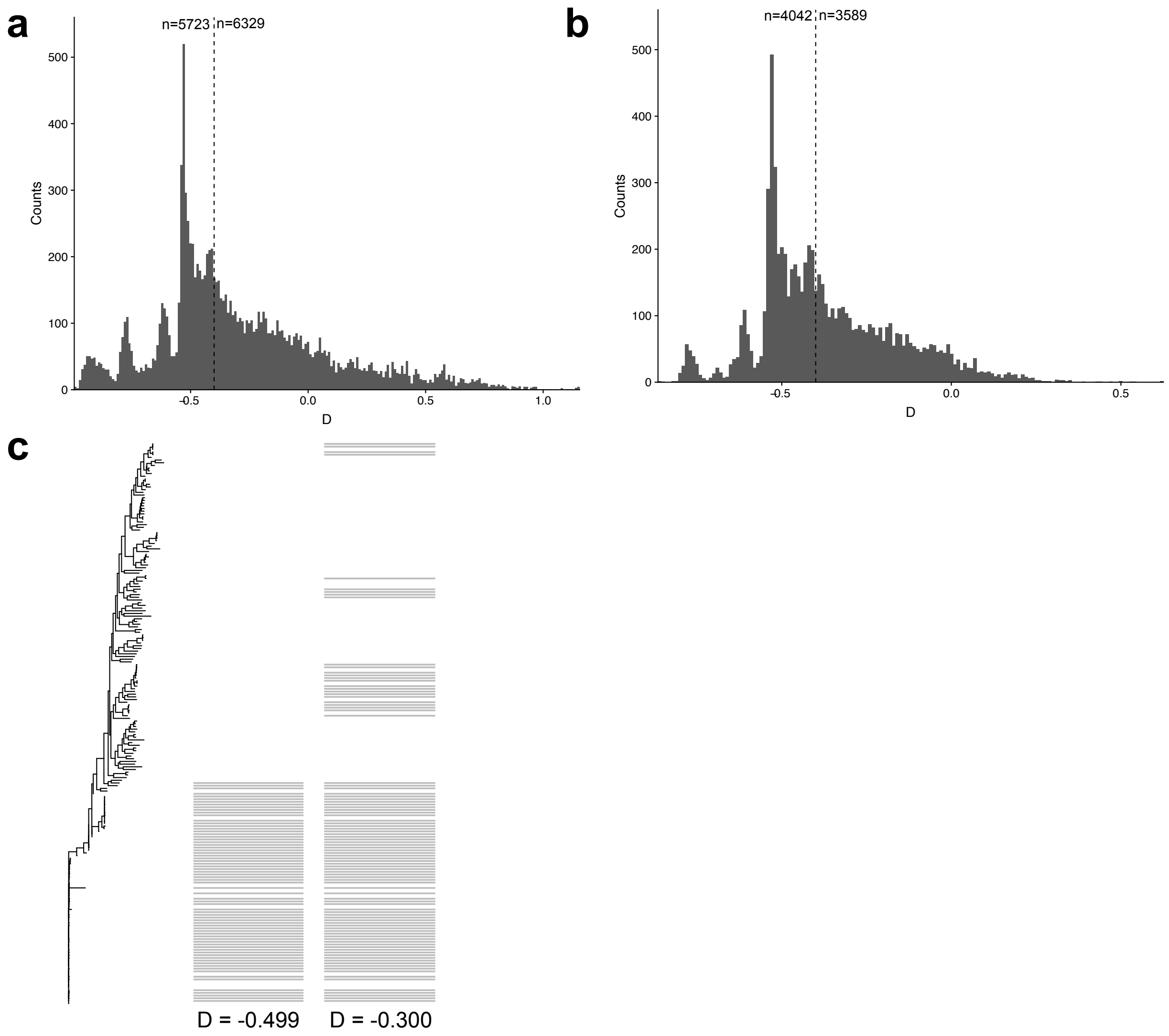


Supplementary Figure 3: **D-value distribution across the nodes found to have significant co-occurring (a) and avoidance (b) relationships in the complete genome dataset (n=209).** The D-value indicates the amount of lineage-independence of a particular gene compared to a phylogeny. Based on the counts and distribution of genes across the core gene phylogeny, a D-value cutoff of -0.4 was chosen, resulting in the inclusion of 6329 and 3589 gene clusters (i.e. nodes), respectively, in the co-occurrence and avoidance datasets. **c.** An example of the distribution of 2 genes across the core gene phylogeny with D values on either side of the chosen threshold.

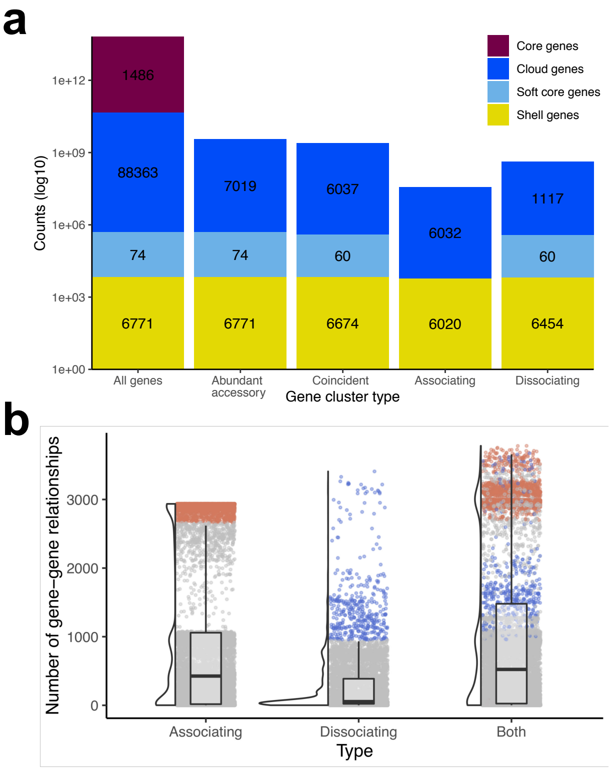


Supplementary Figure 4: **Coincident gene relationships between lineage-dependent and -independent genes. a.** Similar patterns of coincident gene relationships are seen when lineage-dependent genes are included when compared to counts of only lineage-independent genes (**Figure 1c**). **b.** However, the distribution of relationships per gene differs when lineage-dependent genes are included, including an increase in the number of hub genes when compared to those identified between lineage-independent genes (**Figure 1e**). Boxplots display the first and third quartiles, with a horizontal line to indicate the median, and whiskers extend to 1.5 times the interquartile range.

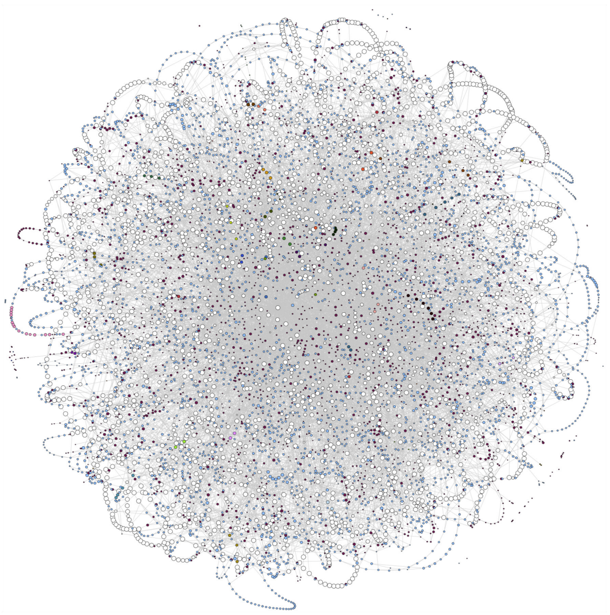

Supplementary Figure 5: **The graph of the *Pseudomonas sp.* pangenome**. The pangenome graph of the 209 *Pseudomonas sp.* genomes as determined by PPanGGOLiN. Each gene cluster (node) is coloured by its connected component with the same colour scheme used in **Figure 1a**. For visibility, cloud genes are not shown.

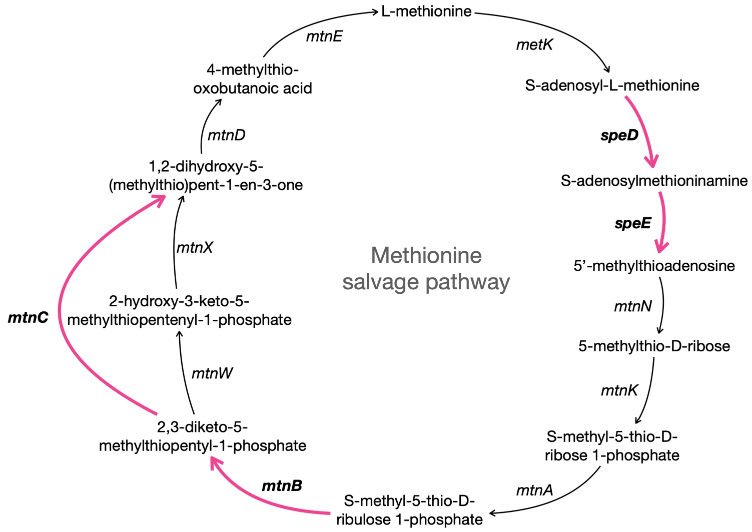


Supplementary Figure 6: **The location of the co-occurring gene set within the simplified pathway of methionine salvage.** Homologs of the gene clusters in question are bolded and the steps they are involved with coloured. These 4 genes were found to significantly co-occur but not all genes within the gene set are co-localized on the genome. This visual of select reactions and metabolites of the methionine salvage pathway is based off of information obtained from KEGG.

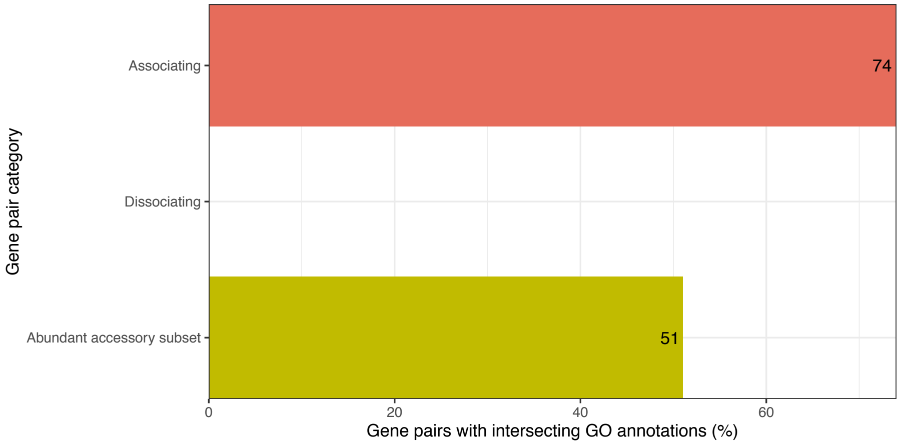


Supplementary Figure 7: **Non-syntenic coincident gene pairs share functionality.** Similar to the analysis of all coincident gene pairs (**Figure 3a**), the results hold when only those with a mean distance of ≥150,000bp between genes are considered. Because dissociating genes by definition are not found in the same genome, this analysis could not be performed on dissociating gene pairs.

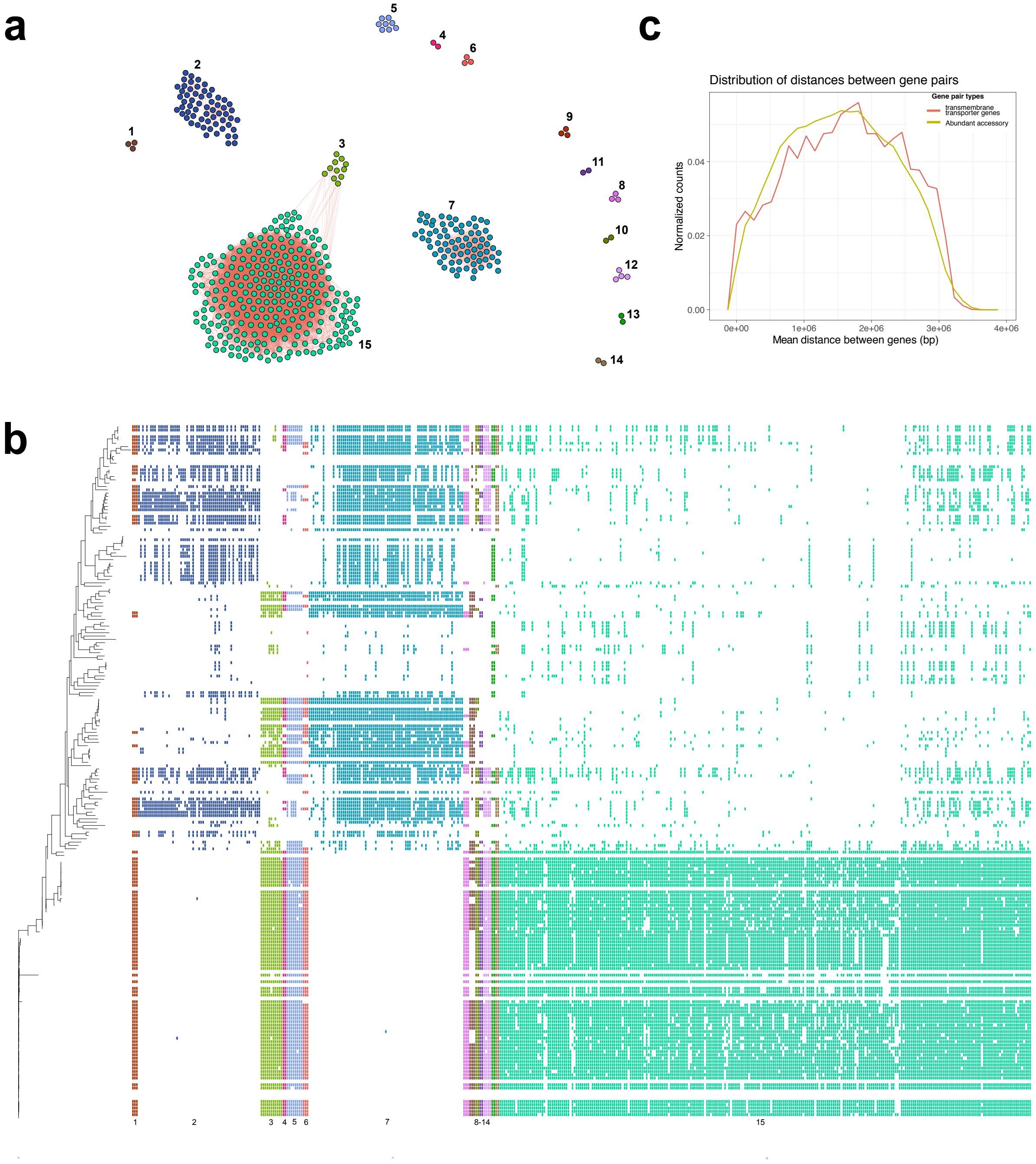


Supplementary Figure 8: **Genes annotated as transmembrane transporters are over represented in both gene association and dissociation relationships. a.** The gene association relationships amongst transmembrane transporters; this network is the same as that displayed in **Figure 3c** but with the dissociation edges hidden for visibility. Labels represent gene cluster numbers. **b.** The mean genomic distance between genes with the GO annotation of transmembrane transporters is evenly distributed across the pangenome. **c.** The presence absence patterns of the gene clusters across the *Pseudomonas sp.* phylogeny. Clusters are coloured and numbered as in part A and **Figure 3c**.

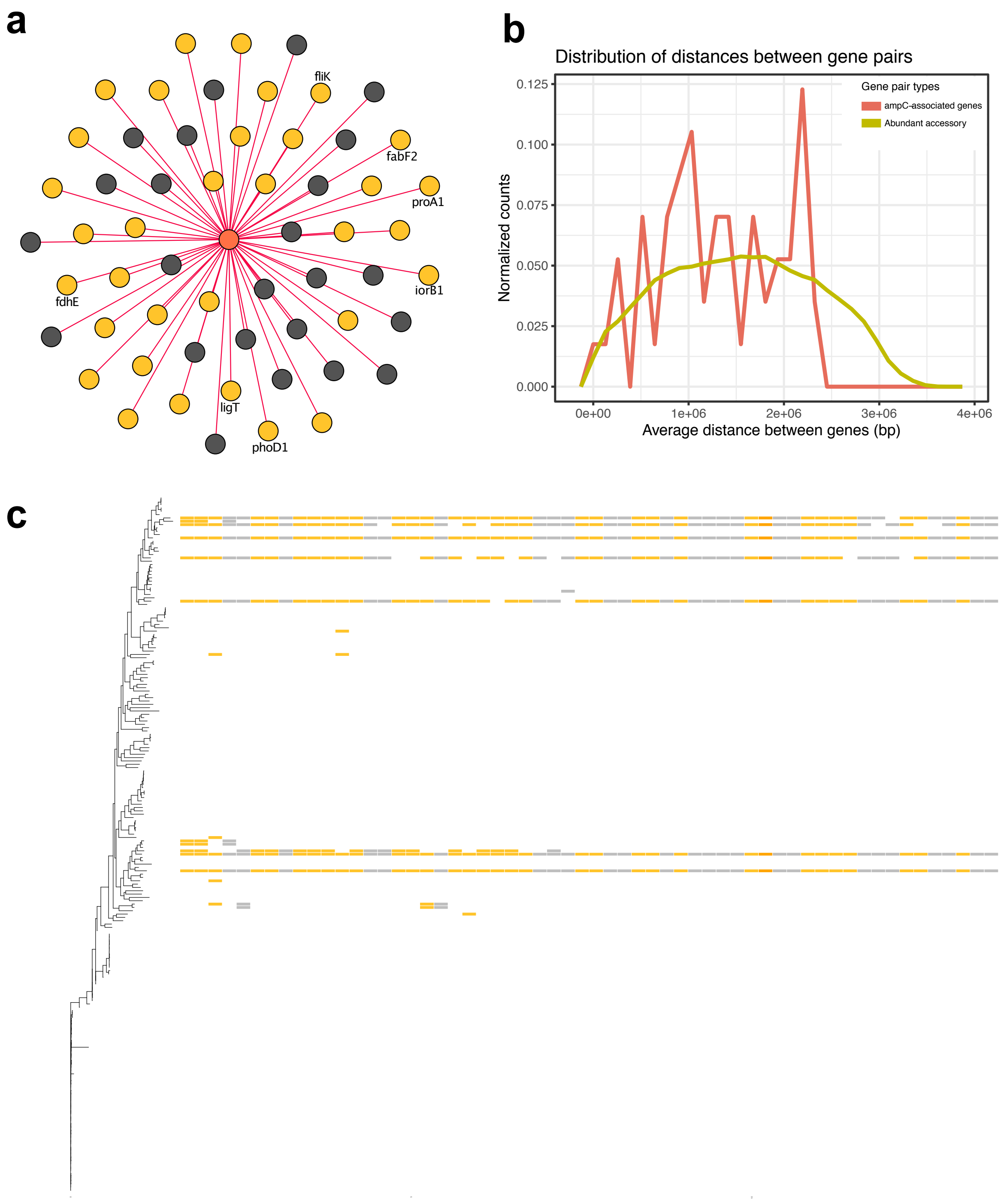


Supplementary Figure 9: **More hypothetical genes co-occur with an *ampC* homolog than would be expected by chance. a.** The network of genes associating with *ampC*. Genes with named annotations are labelled. **b.** These genes do not share significant co-localization with each other when compared with randomly chosen abundant accessory genes. **c.** The presence/absence patterns of these genes across the *Pseudomonas sp.* phylogeny indicates a lack of evolutionary history of these genes. Genes are coloured as to whether they are annotated (**yellow**) or hypothetical (**gray**). The *ampC* homolog is coloured **orange**.

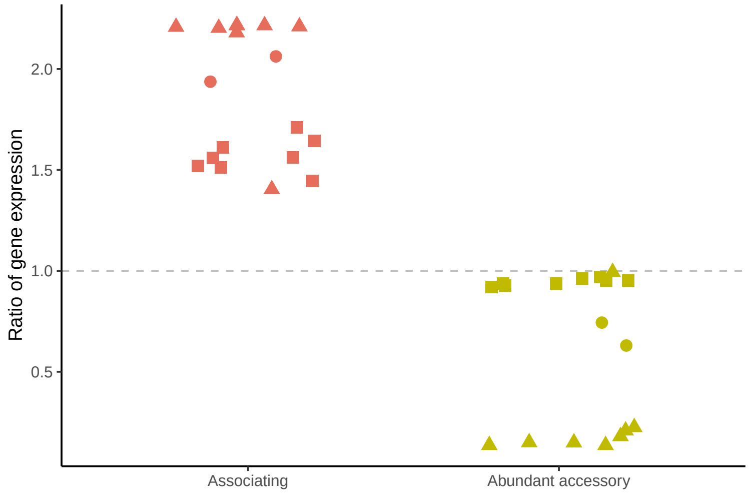


Supplementary Figure 10: **Gene co-expression ratios of non-syntenic gene pairs.** The gene co-expression ratio when only genes with $\geq$ 150,000bp between them were considered.

Supplementary Table 1: **Gene names and descriptions of the gene sets used as examples of high, low, and variable physical distance in Figure 2.** Gene cluster descriptions are based on the descriptions of each gene within the cluster; parentheses indicate how often a given description was observed.

| **Box #** | **Gene cluster ID** | **Gene name/description** |
|---|---|---|
| 1 | group_31493 | hypothetical protein (25); HAD family hydrolase (5); conserved hypothetical protein (1) |
| 1 | group_37342 | nucleotidyltransferase (24); hypothetical protein (2); sugar nucleotidyltransferase (2); putative sugar nucleotidyltransferase (1) |
| 1 | group_49093 | hypothetical protein (26); flagellar assembly protein FliT (3) |
| 1 | group_49094 | flagellar protein FliS (23); flagellar biosynthesis protein FliS (3); B-type flagellar protein FliS (2); hypothetical protein (1) |
| 1 | group_49095 | B-type flagellar hook-associated protein 2 (18); branched-chain alpha-keto acid dehydrogenase subunit E2 (7); flagellar capping protein FliD (4) |
| 1 | group_49096 | hypothetical protein (29) |
| 1 | group_49097 | glycosyl transferase family 1 (14); hypothetical protein (9); flagellar glycosyl transferase, FgtA (3); flagellar glycosyl transferase FgtA (2); glycosyl transferase (1) |
| 1 | group_49098 | hypothetical protein (18); methyltransferase type 12 (10) |
| 1 | group_49099 | flagellar hook-associated protein FlgL (28); flagellar hook-associated protein type 3 FlgL (1) |
| 2 | group_1102 | hypothetical protein (40); dehydrogenase (17); aldose sugar dehydrogenase YliI (9); oxidoreductase (4); putative dehydrogenase (2); aldose sugar dehydrogenase (1); Soluble aldose sugar dehydrogenase, PQQ-dependent (1) |
| 2 | group_1963 | porin (70); Glucose/carbohydrate outer membrane porin OprB precursor (6); Glucose-selective porin OprB (2); carbohydrate porin (1); outer membrane porin OprB (1) |
| 2 | group_7677 | outer membrane protein OprG (66); outer membrane protein W (7); Outer membrane protein OprG precursor (5); Outer membrane protein W precursor (2); OmpW family protein (1) |
| 3 | group_11613 | S-adenosylmethionine decarboxylase proenzyme (66); S-adenosylmethionine decarboxylase (13); S-adenosylmethionine decarboxylase proenzyme, prokaryotic class 1A (1) |

| | | |
|---|---|---|
| 3 | group_6267 | enolase-phosphatase E1 (30); haloacid dehalogenase (23); 2,3-diketo-5-methylthio-1-phosphopentane phosphatase (13); enolase-phosphatase E-1 (4); 2,3-diketo-5-methylthiopentyl-1-phosphate enolase-phosphatase (2); enolase-phosphatase (2) |
| 3 | group_7302 | methylthioribulose-1-phosphate dehydratase (42); methylthioribulose 1-phosphate dehydratase (38); probable sugar aldolase (1) |
| 3 | group_7553 | spermidine synthase (75); polyamine aminopropyltransferase 1 (6) |

Supplementary Table 2: **Summary of statistically enriched GO terms in the association and dissociation datasets.** See SupTable2.xlsx.

Supplementary Table 3: **Summary of genes with transmembrane transporter activity over represented in both association and dissociation gene relationships.** See SupTable3.xlsx.

Supplementary Table 4: **Summary of statistically enriched GO terms with hypothetical pairs.** See SupTable4.xlsx.

Supplementary Table 5: **Summary of genes co-occurring with an *ampC* homolog.** Gene cluster descriptions are based on the descriptions of each gene within the cluster; parentheses indicate how often a given description was observed.

| **Gene cluster ID** | **Gene name/description** |
|---|---|
| group_10590 | hypothetical protein (13) |
| group_1110 | spermidine/putrescine ABC transporter substrate-binding protein PotF (4); putrescine/spermidine ABC transporter substrate-binding protein (3); spermidine/putrescine ABC transporter substrate-binding protein (2); extracellular solute-binding protein (1); polyamine ABC transporter substrate-binding protein (1); putrescine-binding periplasmic protein (1) |
| group_13151 | 2'-5' RNA ligase (17) |
| group_13686 | flagellar hook-length control protein (12); flagellar hook-length control protein FliK (2) |
| group_13726 | hypothetical protein (16) |
| group_13727 | twin-arginine translocation pathway signal protein (16); isoquinoline 1-oxidoreductase subunit beta (1) |
| group_13748 | GNAT family acetyltransferase (10); GNAT family N-acetyltransferase (2); GCN5-like N-acetyltransferase (1); N-acetyltransferase GCN5 (1) |
| group_13811 | hypothetical protein (14) |
| group_13818 | phosphodiesterase (12); phosphodiesterase/alkaline phosphatase D-like protein (2) |
| group_13819 | hypothetical protein (16) |
| group_13820 | hybrid sensor histidine kinase/response regulator (8); histidine kinase (3); multi-sensor hybrid histidine kinase (2); hypothetical protein (1) |
| group_14115 | formate dehydrogenase accessory protein FdhE (13); formate dehydrogenase accessory protein (1) |
| group_14332 | glycosyltransferase (9); glycosyl transferase (3); hypothetical protein (1) |
| group_14389 | aldo/keto reductase (14) |
| group_14455 | peptide transporter (12); amino acid adenylation:thioester reductase (1) |
| group_14473 | chorismate mutase (13) |
| group_15113 | hypothetical protein (13) |
| group_15142 | hypothetical protein (12) |
| group_15152 | hypothetical protein (10); thiolase (2) |
| group_15235 | DUF3077 domain-containing protein (9); hypothetical protein (5) |
| group_15241 | serine protease (11); hypothetical protein (2) |
| group_15274 | hypothetical protein (12) |
| group_15276 | transcriptional regulator CynR (8); transcriptional regulator (4); DNA-binding transcriptional regulator CynR (1) |

| | |
|---|---|
| group_15277 | hypothetical protein (14) |
| group_15280 | ATPase (6); hypothetical protein (6) |
| group_15281 | GCN5 family acetyltransferase (9); GNAT family N-acetyltransferase (2); GCN5-like N-acetyltransferase (1); N-acetyltransferase GCN5 (1) |
| group_15282 | GDP-mannose pyrophosphatase (8); GDP-mannose pyrophosphatase nudK (2); nucleoside diphosphate pyrophosphatase (2); ADP-ribose pyrophosphatase (1) |
| group_15283 | transcriptional regulator (11); Cro/CI family transcriptional regulator (1) |
| group_15284 | hypothetical protein (10); delta-60 repeat domain-containing protein (2) |
| group_15285 | hypothetical protein (11); lipoprotein (1) |
| group_15286 | isochorismatase (10); cysteine hydrolase (1); isochorismatase hydrolase (1) |
| group_15287 | calpastatin (9); hypothetical protein (3) |
| group_15887 | polyketide cyclase (10); MxaD family protein (2); hypothetical protein (1) |
| group_15938 | hypothetical protein (12) |
| group_16075 | methylase (9); hypothetical protein (2); SAM-dependent methyltransferase (1) |
| group_16076 | general secretion pathway protein GspH (10); general secretion pathway protein H (1) |
| group_16077 | hypothetical protein (11) |
| group_16078 | hypothetical protein (9); membrane protein (3) |
| group_16082 | hypothetical protein (9); lipoprotein (2) |
| group_16083 | hypothetical protein (12) |
| group_16084 | hypothetical protein (11) |
| group_16085 | hypothetical protein (12) |
| group_16086 | hypothetical protein (12) |
| group_16087 | NUDIX hydrolase (11) |
| group_16088 | glycerol-3-phosphatase (10); HAD family hydrolase (1); phosphatase (1) |
| group_16091 | acetyltransferase (9); GCN5-like N-acetyltransferase (1); GNAT family N-acetyltransferase (1) |
| group_16092 | class C beta-lactamase (10); beta-lactamase (2) |
| group_16908 | hypothetical protein (11) |
| group_17033 | hypothetical protein (12) |
| group_17040 | hypothetical protein (10); DUF4160 domain-containing protein (1) |
| group_17041 | hypothetical protein (11); lipoprotein (1) |
| group_17048 | glutamate-5-semialdehyde dehydrogenase (12) |
| group_17049 | cupin (6); hypothetical protein (5) |
| group_17050 | glutathione S-transferase (10); glutathione S-transferase-like protein (2) |

| | |
|---|---|
| group_18423 | hypothetical protein (6); endoribonuclease L-PSP (4); endoribonuclease L-PSP family protein (1) |
| group_28806 | (12); tRNA-Arg (3); tRNA-Arg;Dbxref=GeneID:14051617 (1); tRNA-Arg;Dbxref=GeneID:3712881 (1) |
| group_4768 | beta-ketoacyl-[acyl-carrier-protein] synthase II (5); beta-ketoacyl-ACP synthase II (4); 3-oxoacyl-ACP synthase (2); 3-oxoacyl-(acyl carrier protein) synthase II (1) |
| group_6199 | hypothetical protein (11) |
